# Identification of human CD4^+^ T cell populations with distinct antitumor activity

**DOI:** 10.1101/2019.12.31.891317

**Authors:** Michelle H. Nelson, Hannah M. Knochelmann, Stefanie R. Bailey, Logan W. Huff, Jacob S. Bowers, Kinga Majchrzak, Megan M. Wyatt, Mark P. Rubinstein, Shikhar Mehrotra, Michael I. Nishimura, Kent E. Armeson, Paul G. Giresi, Michael J. Zilliox, Hal E. Broxmeyer, Chrystal M. Paulos

**Affiliations:** Department of Microbiology & Immunology, Medical University of South Carolina, Charleston, SC, USA; Department of Dermatology & Dermatologic Surgery, Medical University of South Carolina, SC, USA; Department of Surgery, Medical University of South Carolina, Charleston, SC, USA; Department of Surgery, Stritch School of Medicine, Loyola University Chicago, Maywood, IL, USA; Department of Public Health Sciences, Medical University of South Carolina, Charleston, SC, USA; Epinomics, Menlo Park, CA, USA; Department of Public Health Sciences, Stritch School of Medicine, Loyola University Chicago, Maywood, IL, USA; Department of Microbiology & Immunology, Indiana University School of Medicine, Indianapolis, IN, USA

**Author notes:** Contributed equally to this work (Michelle H. Nelson, Hannah M. Knochelmann). Corresponding author: Chrystal M. Paulos, Hollings Cancer Center, HO606B; MSC 509, Medical University of South Carolina, 86 Jonathan Lucas St, Charleston, SC 29425.

## Abstract

How naturally arising human CD4^+^ T helper subsets impact tumor immunity is unknown. We reported that human CD4^+^CD26^high^ T cells elicit potent immunity against solid tumor malignancies. As CD26^high^ T cells secrete type-17 cytokines and have been categorized as Th17 cells, we posited these helper populations would possess similar molecular properties. Herein, we reveal that CD26^high^ T cells are epigenetically and transcriptionally distinct from Th17 cells. Of clinical significance, CD26^high^ T cells engineered with a chimeric antigen receptor (CAR) ablated large human tumors to a greater extent than enriched Th17, Th1, or Th2 cells. Moreover, CD26^high^ T cells mediated curative responses in mice, even when redirected with a suboptimal CAR and without the aid of CD8^+^ CAR T cells. CD26^high^ T cells co-secreted effector cytokines at heightened levels and robustly persisted. Collectively, our work reveals the potential of human CD4^+^ T cell populations to improve durability of solid tumor therapies.

## Introduction

We previously reported that CD26 distinguishes three human CD4^+^ T cell subsets with varying degrees of responsiveness to human tumors: one with regulatory properties (CD26^neg^), one with a naive phenotype (CD26^int^), and one with a durable stem memory profile (CD26^high^) (*1*). Adoptively transferred tumor-specific CD26^high^ T cells persisted and regressed difficult-to-treat malignancies superior to CD26^neg^ T cells and surprisingly, slightly better than naive CD26^int^ T cells. CD26^high^ T cells secreted Th17 cytokines, including IL-17A, TNFα and IL-22. These previous findings reveal that CD26^high^ T cells are promising for immunotherapies.

The therapeutic potency of Th17 cells over Th1 or Th2 cells has been reported by many groups using mouse model systems (*2*–*5*). Surprisingly, the impact of tumor-specific *human* Th17 cells have not been fully explored in the context of adoptive T cell transfer (ACT) therapy for cancer. Given the sizeable expression of master transcription factor RORγt and production of IL-17 by human CD26^high^ T cells (1), we postulated that these cells would eradicate tumors to the same extent as classic Th17 cells (i.e. CCR4^+^CCR6^+^CD4^+^) when redirected with a chimeric antigen receptor (CAR) and infused into hosts bearing human tumors (*6*). Moreover, the therapeutic potential of naturally arising human CD4 subsets—sorted from the peripheral blood via classic surface markers—engineered with a CAR has yet to be elucidated.

We report herein that CD26^high^ T cells are molecularly and functionally distinct from Th17 cells. CD26^high^ T cells are robustly therapeutic compared to Th17, as demonstrated by their capacity to persist and eradicate large human tumors in mice. Additional investigation uncovered that CD26^high^ T cells were more effective than Th1 or Th2 cells as well. We found that the molecular and epigenetic properties of CD26^high^ T cells are distinct from Th17 cells, which might support their persistence and sustained responses to large tumors.

## Results

### CD26^high^ T cells possess a dynamic cytokine profile

We reported that CD4^+^ T cells expressing high CD26 levels (termed CD26^high^ T cells) secrete IL-17A and elicit potent tumor immunity when redirected with a CAR compared to sorted CD26^int^ or CD26^low^ T cells (*1*). While CD26^high^ T cells are categorized as Th17 cells, the functional profile of sorted human CD26^high^ T cells compared to classic Th17 cells as well as other known helper subsets has never been tested. Given the abundance of IL-17 produced by CD26^high^ T cells, we suspected that they would possess a similar cytokine profile as classic Th17 cells. To first address this question, we measured the level and type of cytokines produced by various CD4^+^ subsets, which were sorted from the peripheral blood of healthy individuals via extracellular markers (Figure 1A) (*6*). This sort yielded Th1 (CXCR3^+^CCR4^−^CCR6^−^), Th2 (CXCR3^−^CCR4^+^CCR6^−^), Th17 (CCR4^+^CCR6^+^CXCR3^+/−^) and CD26^high^ T cells with high purity (>90%). As expected, Th1 cells expressed CXCR3, Th2 cells expressed CCR4, and Th17 cells expressed CCR4 and CCR6. CD26^high^ cells expressed high CXCR3 and CCR6 but nominal CCR4 on their surface (Figure 1B).

**Figure 1.**
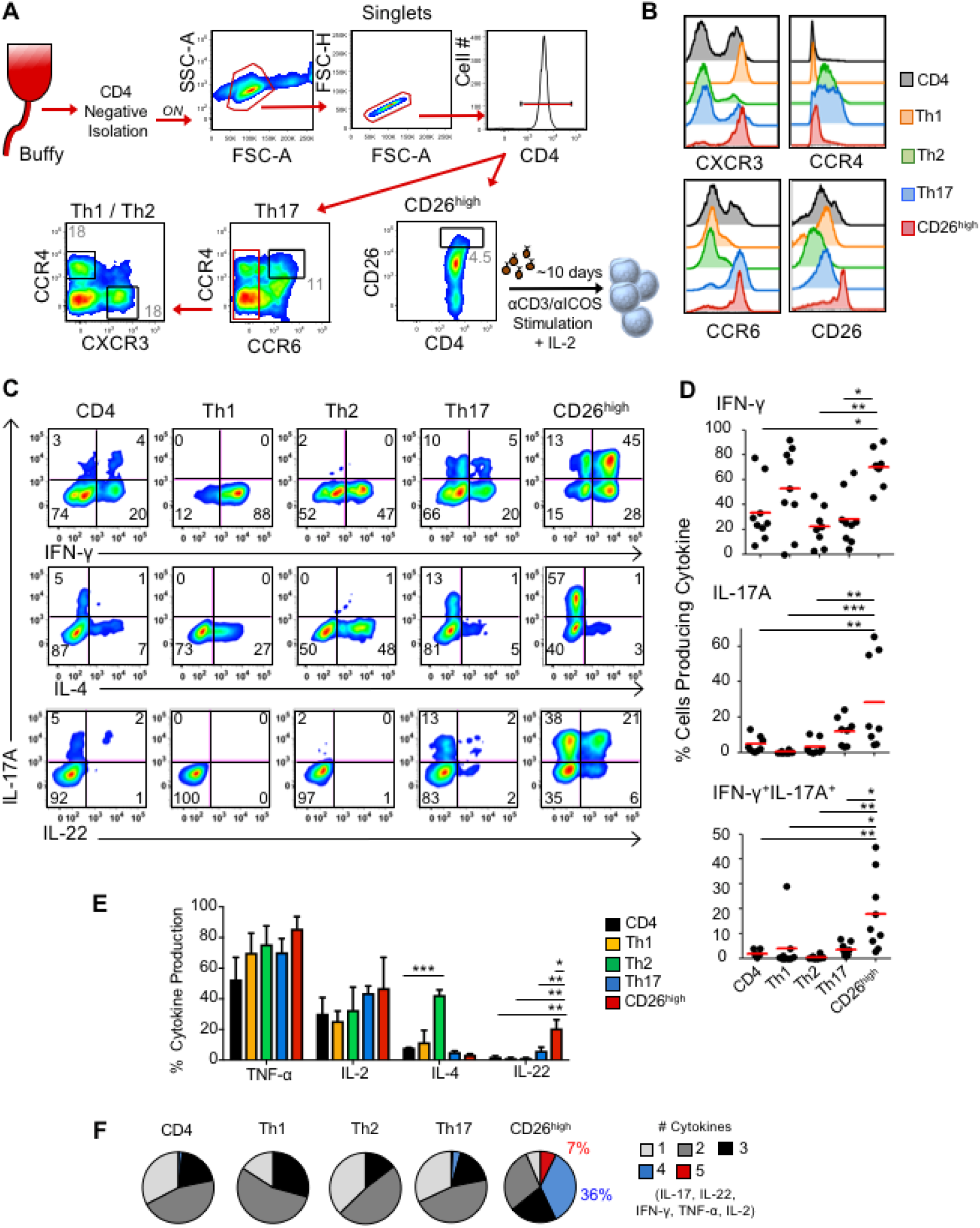
CD4^+^CD26^high^ T cells possess a dynamic cytokine profile. **A**) CD4^+^ subset sorting scheme. CD4^+^ lymphocytes were negatively isolated using magnetic beads from normal donor PBL. Th17 cells were sorted from CCR6^+^CCR4^+^ gate. Th1 and Th2 cells are both CCR6^−^ and subsequently sorted via CXCR3 or CCR4, respectively. CD26^high^ cells were sorted independently based on CD26 expression. **B**) Chemokine receptor profile post sort. **C**) CD4^+^ T cell subsets were stimulated with αCD3/ICOS beads at a ratio of 1 bead:10 T cells and expanded in IL-2 (100IU/ml). Ten days following activation, the 5 different cell subsets were examined for their intracellular cytokine production. Dot plot representation of IL-17, IFN-γ, IL-4, and IL-22 expression by flow cytometry. **D**) Graphical representation of at least 8 normal donors from independent experiments demonstrating IFN-γ and IL-17 single and double producing cells by flow cytometry. **E**) Graphical representation of 10 normal donors demonstrating cytokine-producing cells by flow cytometry. 2-3 replicates each. Compared to CD26^high^ *, *P*<0.05; **, *P*<0.01; ***, *P*<0.001; ANOVA, Tukey post-hoc comparisons. **F**) Cells were gated on cytokine-producing cells to quantify cells that produced between one and five cytokines simultaneously. Cytokines of interest were IL-17, IFN-γ, IL-2, IL-22 and TNF-α. Representative of 5 experiments.

CD26^high^ T cells were not restricted to a Th17-like functional profile (Figure 1C). Instead, CD26^high^ T cells secreted more IL-17A (58 vs. 15%), IL-22 (27 vs. 4%) and IFN-γ (73 vs. 25%) than Th17 cells. CD26^high^ T cells produced nearly as much IFN-γ (73 vs. 88%) as Th1 cells but far less IL-4 (3 vs 29%) than Th2 cells. We consistently observed this functional pattern in CD26^high^ T cells from several healthy individuals (Figure 1C-E). On a per-cell basis, CD26^high^ T cells concomitantly secreted 4 (35%) to 5 (7%) cytokines, a dynamic process not manifested in other subsets (Figure 1F). Collectively, our data suggest that CD26^high^ T cells have a distinct functional profile from classic Th17 cells.

### CD26^high^ T cells display a unique chromatin landscape

Given the functional profile of CD26^high^ T cells, we hypothesized that the epigenetic landscape of these cells at resting state would be different than Th17 cells. To test this idea, we sorted naïve, Th1, Th2, Th17 and CD26^high^ T cells from the blood of 5 different healthy donors and profiled their chromatin accessibility with Assay for Transposase-Accessible Chromatin with high-throughput sequencing (ATAC-seq). CD26^high^ T cells contained peaks displaying enhancer accessible regions near various transcription factors (TF) known to direct Th1 (such as *Tbx21* and *EOMES*) and Th17 (*RORC*) cell lineage development while displaying suppressor regions near TF genes known to regulate Th2 development, such as *GATA3* (Figure 2A-B). While *Tbx21* and *EOMES* were more accessible in both Th1 and CD26^high^ T cells, they were repressed in naïve, Th2 and Th17 cells (Figure 2B). Moreover, a core of other accessible regions in Th1-related TFs, such as *MGA, STAT2, STAT1* and *STAT5A*, were pronounced in Th1 and CD26^high^ T cells (Figure 2A). As expected, accessible regions surrounding *GATA3* were enhanced in Th2 cells and interestingly in Th17 cells. Other enhancer accessible regions surrounding Th2-like TFs, such as *GATA1, GATA2, GATA4, GATA5, GATA6, PAX4, YY1, PITX2* and *GFI1* were distinguished in Th2 and Th17 cells (Figure 2A). Similar to Th17 cells, chromatin accessible regions near the *RORA, RORB*, and *STAT3* loci were enhanced in CD26^high^ T cells but suppressed in naïve, Th1 and Th2 cells (Figure 2A-B). Th1 cells more closely aligned with the epigenetic landscape of naïve cells, as they both expressed accessible chromatin regions neighboring TFs in the stem and development pathways, including *TCF1, LEF1, CTCF, DNMT1* and *ZFP161* (Figure 2A). Yet, certain accessible regions in naïve cells were also heightened in both CD26^high^ and Th1 subsets, including *STAT1, STAT2, IRF1, IRF2, IRF3, IRF5, IRF7, IRF8* and *ZNF683* (Figure 2A).

**Figure 2.**
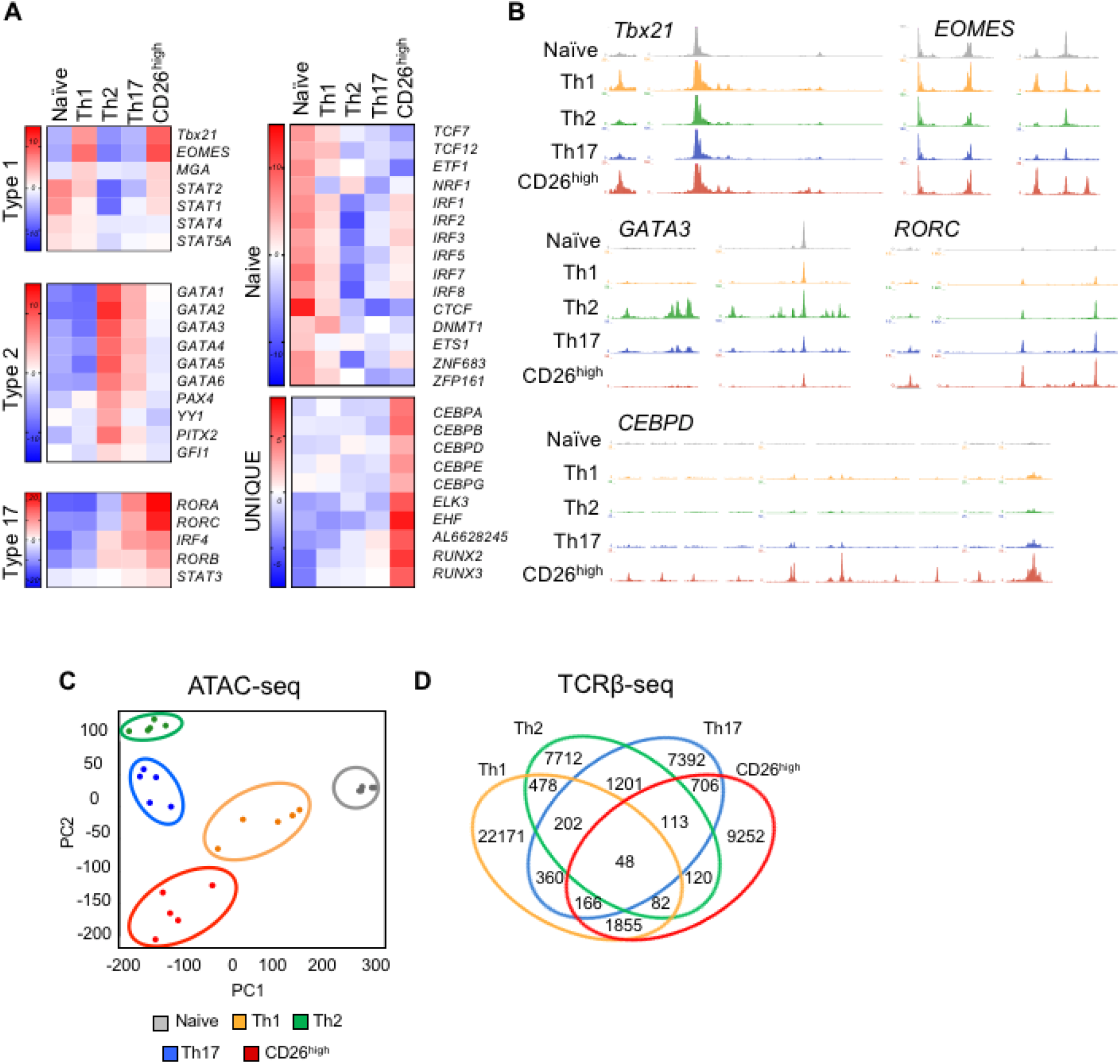
The epigenetic and molecular signature of CD26^high^ T cells are unique. **A**) Assay for Transposase-Accessible Chromatin sequencing (ATAC-seq) analysis describing chromatin accessibility in FACS sorted CD4^+^ subsets (Naïve, Th1, Th2, Th17, CD26^high^) organized by transcription factor networks known to describe Th1, Th2, Th17 and naïve subsets. Accessible transcription regions unique to CD26^high^ T cells are also shown. Compiled from 5 healthy donors. **B**) UCSC genome browser tracks for sorted CD4^+^ subsets around classical T helper transcription factors from ATAC-seq analysis. **C**) ATAC-seq principal component analysis of sorted T cell subsets analyzed at resting state. n=5 donors. **D**) TCRβ sequencing of CD26^high^, Th17, and Th1 cells sorted from peripheral blood of healthy donors demonstrates unique or shared clonotypes. Venn diagram illustrates percentage of unique or shared TCRβ sequences. The relative frequencies (standardized to sum to 1.0): CD26^high^ only = 0.237, Th1 only = 0.487, Th17 only = 0.196, CD26^high^ & Th1 = 0.041, CD26^high^ & Th17 = 0.020, Th1 & Th17 = 0.015, All three = 0.004, log-linear model.

Despite overlap with Th1 and Th17 cells, CD26^high^ T cells possessed a unique set of differentially accessible elements relative to other subsets. Open accessible regions in the CCAAT/enhancer-binding protein family (C/EBP), which function as TFs in processes including cell differentiation, motility and metabolism, were among the most unique and differentially expressed in CD26^high^ T cells (Figure 2A-B). Along with *CEBPs*, *ELK3*, important for cell migration and invasion, and *RUNX*, which promotes memory cell formation, were enhanced in CD26^high^ T cells. Principal component analysis of the genome-wide open chromatin landscape of these 25 samples showed that CD26^high^ T cells cluster separately from naïve, Th1, Th2 and Th17 cells (Figure 2C). We verified the distinct characteristics of CD26^high^ versus Th17 cells using gene array (Figure S1A-B). Further, as helper subsets have been reported to express a particular TCRβ repertoire (*7*), we defined the frequency and likelihood of TCRβ clonotype overlap between various sorted subsets and found nominal overlap between CD26^high^ cells and other helper subsets (Figure 2D & Figure S1C). Collectively, we conclude that the epigenetic landscape and TCR repertoire of CD26^high^ cells differs substantially from that of classic CD4^+^ subsets.

### Single cell sequencing reveals that CD26^high^ T cells are molecularly unique from Th17 cells

Single-cell transcriptome analysis also supported that CD26^high^ T cells are distinguished from Th17 cells based on differential clustering from that of bulk CD4^+^ and Th17 cells (Figure 3A). Interestingly, a cluster of Treg-like cells was present within the sorted Th17 population (Figure 3B), as demonstrated by heightened *FOXP3*, *IL2RA*, and *TIGIT* and reduced *IL7R* transcript, but was not found within CD26^high^ T cells. A small cluster of Th1-like cells was identified within the sorted bulk CD4^+^ population, as indicated by elevated *TBX21, GZMH, PRF1, CCL5* and *CXCR3* but nominal transcripts associated with Th17 or Treg cells, such as *CCR6, CCR4, RORC* and *FOXP3*. Transcripts describing naïve-like cells including *SELL*, *CCR7, CD27*, and *LEF1* were expressed at slightly higher levels in bulk CD4^+^ cells than other populations. In concurrence with their chromatin accessibility, *CEBPD* transcripts were elevated in CD26^high^ T cells compared to bulk CD4^+^ or Th17 cells, potentially indicating a bioenergetic profile resistant to oxidative stress (*8*). Taken together, these data suggest that CD26^high^ T cells are unique from Th17 cells, yet their relative clinical potential in cancer immunotherapy remained unknown.

**Figure 3:**
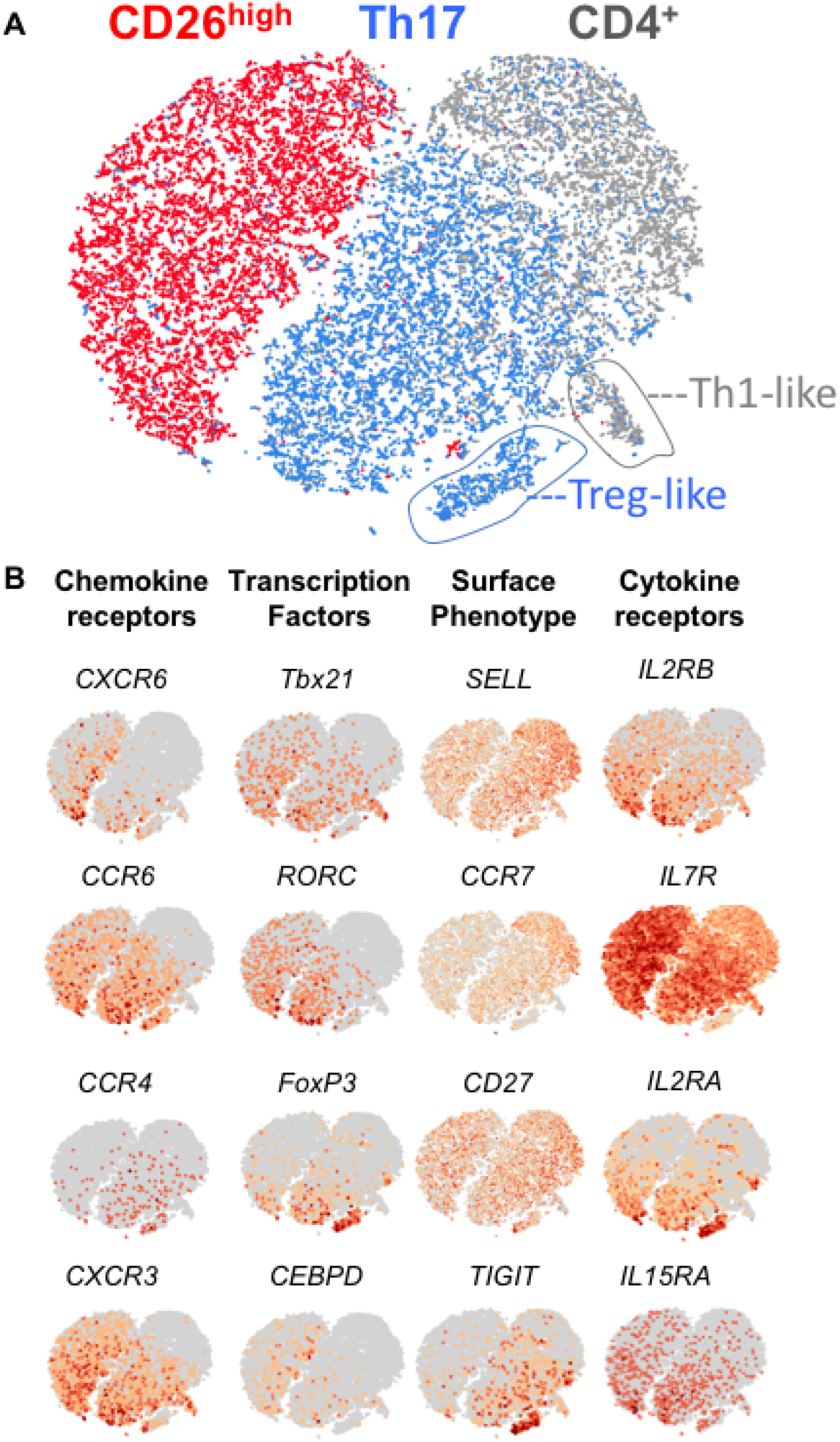
CD4^+^CD26^high^ T cells are distinguished from Th17 cells via single-cell sequencing. Total CD4^+^, CD26^high^ and Th17 cell subsets were sorted from the peripheral blood of healthy donors and ~3000 cells assayed by single cell RNA sequencing. **A**) Data were analyzed by t-Distributed Stochastic Neighbor Embedding (t-SNE). **B**) t-SNE plot overlaid with mRNA expression of chemokine receptors, transcription factors, memory markers and cytokine receptors. Representative of 3 healthy donors.

### CD26^high^ T cells lyse tumor target cells *in vitro*

Given the pronounced capacity of CD26^high^ T cells to co-secrete multiple cytokines, we tested if they would be more effective at lysing human tumors than Th1, Th2 or Th17 cells *in vitro*. To address this, we engineered these helper populations to express chimeric antigen receptor that recognizes mesothelin (meso-CAR) and co-cultured them with mesothelin-positive K562 tumor cells (Figure S2A). As anticipated, CD26^high^ T cells lysed tumor targets at a lower effector to target (E:T) ratio compared to all other subsets when co-cultured overnight (Figure S2B). In this assay, Th1, Th17 and bulk CD4^+^ T cells similarly lysed targets at equal E:T ratios, whereas a greater number of Th2 cells were needed to lyse targets. Finally, after co-culture with target cells, CD26^high^ T cells produced as much IFN-γ and IL-17 as Th1 and Th17 cells, respectively (Figure S2C). Thus, CD26^high^ T cells are highly polyfunctional and mount robust responses against tumors *in vitro*.

### CD26^high^ T cells demonstrate enhanced tumor immunity compared to other helper subsets

Next, we set out to test the relative antitumor activity of these CD4^+^ T helper populations *in vivo*. As in Figure S2A, we engineered sorted Th1, Th2, Th17, CD26^high^, and bulk CD4^+^ T cells to express mesothelioma specific-CAR and infused them into NSG mice bearing a large established tumor. Note that we used a 1^st^ generation meso-CAR, reported by our colleagues to be less therapeutic than 2^nd^ generation meso-CARs (*9*), as we surmised this approach would generate a treatment window to address whether CD26^high^ T cells lyse tumor to a greater extent than other subsets. CD8^+^ T cells (10 day-expanded) were also redirected with this 1^st^-generation CAR and co-infused with these various CAR-CD4^+^ subsets (Figure 4A). CD26^high^ T cells eradicated tumors while Th17 cells only regressed tumors short term (Figure 4B-C). Th17 cells were more effective than Th1 or bulk CD4^+^ T cells at transiently clearing tumors, while Th2 cells were the least effective (Figure 4B-C). Ultimately, mice treated with CD26^high^ T cells survived significantly longer (Figure 4D), which was associated with higher CD4^+^ and CD8^+^ CAR T cell persistence compared to other helper subsets (Figure 4E). Moreover, co-transferred CD4^+^CD26^high^ cells improved the function of CD8^+^ CAR-engineered T cells, as both persistence of CD8^+^ IFN-γ^+^ and CD8^+^ IFN-γ^+^/IL-2^+^/TNF-α^+^ CAR T were heightened in the spleen (Figure S3A-C). These findings suggested that CD4^+^CD26^high^ CAR T cells persisted and promoted the function of co-transferred CD8^+^ CAR T cells.

**Figure 4.**
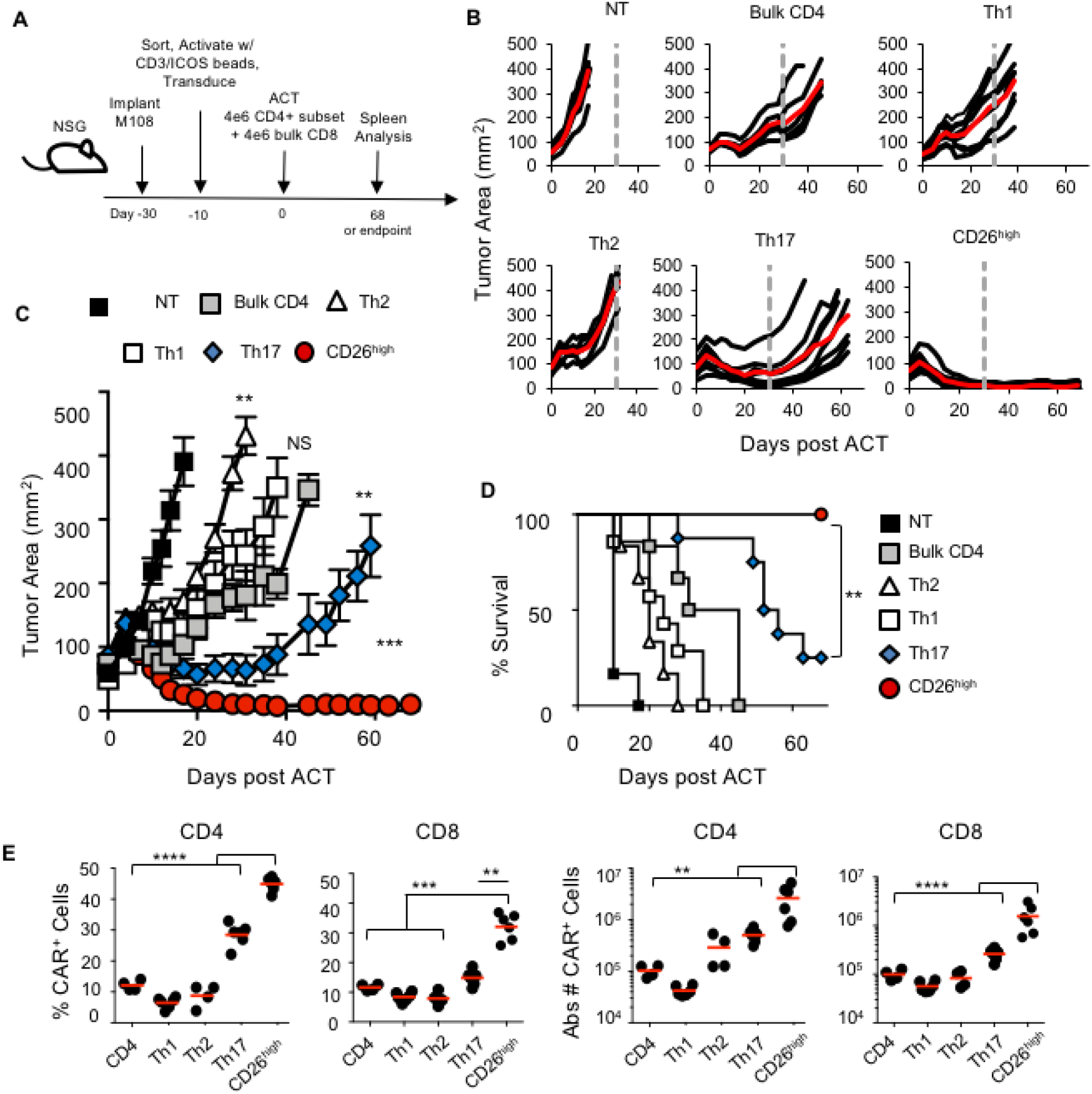
Human CD26^high^ T cells ablate large human tumors and persist relative to other CD4^+^ T cell subsets. **A**) ACT schematic. Th1 (CXCR3^+^), Th2 (CCR4^+^), Th17 (CCR4^+^/CCR6^+^), CD26^high^ or bulk CD4^+^ cells were sorted from normal donor PBL and expanded with αCD3/ICOS bead at a 1 bead:10 T cell ratio. Cells were transduced with a 1^st^ generation mesothelin-specific CD3ζ CAR and expanded with IL-2. NSG mice bearing mesothelioma were treated with 4×10^6^ transduced, sorted CD4^+^ cells + 4×10^6^ transduced CD8^+^ cells and 50,000 IU IL-2 was given to each mouse daily for 3 days. **B**) Single tumor curves overlaid with average curve (red) and **C**) average tumor curves of 6-9 mice/group. All groups were significantly different from NT, *P* < 0.005. CD4 vs. Th1 NS; CD4 vs. Th2, *P =* 0.0015 (**); CD4 vs. Th17, *P =* 0.0035 (**); CD4 vs. CD26^high^, *P =* 0.0003 (***); Th17 vs. CD26^high^, *P* = 0.008 (**); polynomial regression. **D**) The percentage of mice surviving with tumor size below the 200mm^2^ threshold. **E**) Spleens were analyzed by flow for the percentage and total number of CD3^+^CAR^+^CD4^+^ or CD8^+^ cells at day 68 (Th17 and CD26^high^) or group endpoint (CD4, Th1, Th2). n=4-6 mice/group. Compared to CD26^high^ **, *P* < 0.01; ***, *P* < 0.001; ****, *P* < 0.0001; ANOVA, Tukey post-hoc comparisons.

We sought to uncover if CD8^+^ T cells partnered with CD26^high^ T cells to mediate the long-term survival in mice administered this therapy. Given the polyfunctionality of CD26^high^ cells *in vitro*, we posited that CD26^high^ CAR T cells may not require CD8^+^ CAR T cells for productive immunity. To address this question, we transferred CD4^+^CD26^high^ CAR T cells with or without CD8^+^ CAR T cells into NSG mice bearing M108 tumors (Figure S4A-B). Indeed, CD4^+^CD26^high^ CAR T cells did not require the presence of CD8^+^ CAR T cells to regress tumors, and CD8^+^ CAR T cells alone were not therapeutic long term (Figure S4B).

Finally, we questioned whether the CAR signaling in CD4^+^CD26^high^ cells was critical to improve persistence of CD8^+^ CAR T cells in the tumor, or whether their presence alone (i.e. redirected with a non-signaling CAR) could support CD8^+^ CAR T cells. To address this question, CD8^+^ and CD26^high^ T cells were redirected with either a full-length signaling meso-CAR-ζ or a truncated TCR-ζ domain without signaling capability (Δζ) but could still recognize mesothelin and analyzed their presence in tumors. We found that meso-ζ-CD26^high^ cells, either co-infused with meso-Δζ-CD8^+^ or with meso-ζ-CD8^+^ T cells, promoted CD45^+^ immune infiltration in M108 tumors 84 days post adoptive transfer (Figure S4C). Conversely, CAR T cells did not persist if transferred with meso-Δζ-CD26^high^ cells. Collectively, our work reveals that meso-CAR CD4^+^CD26^high^ cells are cytotoxic *in vitro* and *in vivo*, regress tumors in the absence of CD8^+^ T cells and require tumor-reactive CD3ζ signaling to persist.

## Discussion

CAR T cells are therapeutic in many patients with hematological malignancies but have been less effective thus far against solid tumors, owing in part to the oppressive tumor microenvironment and poor persistence. Many efforts for overcoming these obstacles include modulating T cell trafficking, targeting, cytokine delivery, co-stimulation, and improving cell persistence among other strategies reviewed previously (*10*). CD4^+^ T cells help cytotoxic CD8^+^ T cells and when CAR engineered, have the ability to improve longevity of responses against hematological malignancies (*11, 12*). Here we reveal that naturally arising CD4^+^ T cell subsets in the peripheral blood differentially impact efficacy of CAR T cell therapy. For the first time, we demonstrate that CD4^+^CD26^high^ T cells redirected with CAR possess enhanced functional and antitumor activity versus classic subsets (Th1, Th2, Th17) or unselected CD4^+^ T cells, as summarized visually in Figure 5.

**Figure 5.**
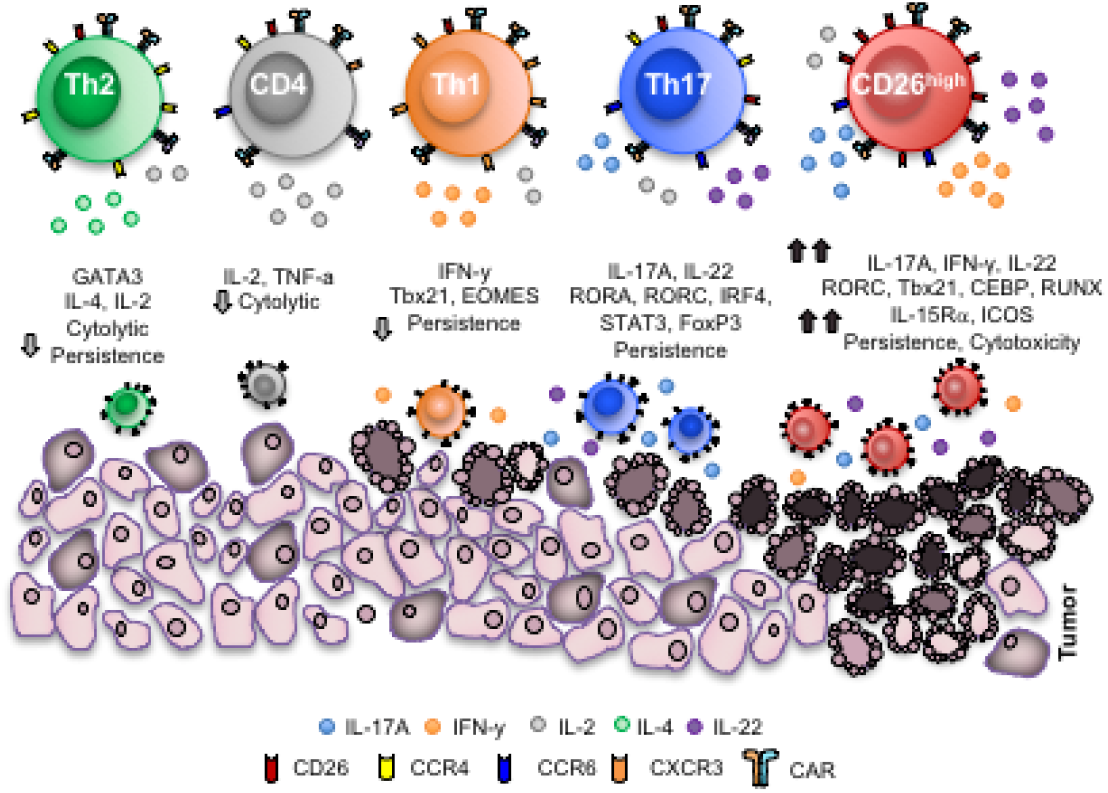
CD4^+^CD26^high^ T cells possess distinct antitumor and molecular properties relative to other helper subsets. CD26^high^ T cells have been described herein for use in adoptive T cell transfer therapy. These cells produce heightened levels of cytokines including IL-17, IFN-γ, IL-22, IL-2 and can co-secrete these cytokines. CD26^high^ T cells have a distinct chromatin landscape with accessible regions near *RORC*, *Tbx21*, *CEBP*, and *RUNX* transcription factors, and have a unique transcriptional signature. These cells are cytotoxic, multi-functional and inflammatory. Overall, CD26^high^ T cells persist and regress tumors to a remarkably greater extent than other CD4^+^ T cells *in vivo* and represent a novel CD4^+^ helper population with potent antitumor properties.

CD26^high^ T cells derived from the peripheral blood of healthy individuals were polyfunctional, co-secreting elevated IL-17A, IFN-γ and IL-22 while classic Th1, Th2 or Th17 cells lacked this dynamic functional profile. Moreover, CD26^high^ T cells have unique epigenetic and molecular properties versus Th17 cells. As well, their TCRβ repertoire does not overlap profoundly with other helper subsets. Of clinical significance, CD4^+^CD26^high^ T cells persisted long-term and ablated mesothelioma in mice when *ex vivo* engineered with CAR unlike bulk CD4^+^, Th1, Th2, or Th17 cells. These cells could improve persistence of co-transferred CD8^+^ CAR T cells yet did not require CD8^+^ T cells for tumor regression. Notably, sorting Th17 cells by CCR4^+^CCR6^+^ yielded an IL-17^+^ population also containing FoxP3^+^IL-2Rα^high^ Tregs, which was not present when sorting CD26^high^ T cells. This work could yield future insight into new methods of sorting T cells to improve CAR therapies by generating more functional T cells.

CD26 has many properties that could impact T cell immunity and our work supports this concept (*13*). CD26 regulates distinct T cell functions, including: a) enzymatic cleavage of chemokines that regulate migration (*14*); b) induction of CD86 on APC via CD26/Caveolin-1 co-stimulation, in turn activating T cells (*15, 16*); c) conversion of adenosine (in tumors) to non-suppressive inosine via docking adenosine deaminase (*17, 18*) and d) binding extracellular matrix proteins (*19*), which may help CD26^high^ T cells infiltrate and remain in tumor. CD26 expressing cells further have high levels of chemokine receptors on their cell surface including CCR2 which promote their recruitment and migration capability and are associated with rapid functional recall responses (*1, 20*). CD26^+^ T cells have been associated with exacerbating various autoimmune manifestations, including rheumatoid arthritis (RA) (*21*), multiple sclerosis (MS) (*22*), graft versus host disease (GVHD) (*23, 24*) and diabetes (*25*). Conversely, levels of CD26 enzymatic activity and the number of CD26^+^ T cells decrease in the blood of melanoma patients as their disease progresses (*26*). This clinical data might suggest that CD26 itself plays a role in augmenting T cell-mediated tumor immunity. Indeed, the CD26 molecule possesses many functions that could be attributed to enhanced antitumor responses, but which one(s), if any, have not yet been elucidated.

It remains possible that none of the many CD26 properties are responsible for regulating the remarkable antitumor activity of these cells. Rather, high CD26 expression may mark lymphocytes with durable persistence. CD26^high^ T cells may have a competitive advantage in the tumor compared to other lymphocyte populations due to their function or perhaps resistance to oxidative stress within the tumor microenvironment suggested by open chromatin and heightened transcription of *CEBPD*. Study of the importance of *CEBPD* to CD26^high^ T cell immunity is underway in our lab. Finally, while our work shows that enriching T cell subsets can improve sub-optimal CAR constructs lacking costimulation, it will be important to clinically elucidate the impact of costimulatory domains on persistence and durability of CD4^+^ T cell populations. Future investigation of the unique CD26^high^ T cell signature discovered herein will reveal the importance of these characteristics to T cell function and provide novel approaches to enhance tumor immunity.

There are many implications from our findings given the significant antitumor responses mediated by CD26^high^ T cells in a mouse model of large established human mesothelioma. The epigenetic and molecular landscape of these helper subsets will permit investigators to address novel questions regarding their function in the immune system. Future work to translate, target and redirect these cells to eradicate tumors or target cells inducing autoimmunity in the clinic could provide new treatment options for a vast array of diseases. Clinical trials are now underway based on our findings to evaluate the potential of CD4^+^CD26^high^ T cells in patients.

## Methods

### Study Design

*Sample Size:* As these experiments were exploratory, there was no estimation to base the effective sample size; therefore, we based our animal studies using sample sizes ≥ 5. *Rules for Stopping Data Collection:* experimental endpoints were designated prior to study execution. Tumor control studies were conducted over ~70 days. *Data Inclusion/exclusion:* for experiments reported herein, animals were only excluded if tumors were very small or not measurable, a rule established prospectively prior to any therapy initiation. *Outliers:* Outliers were reported. *Endpoints:* Tumor endpoint was reached when tumor area exceeded 400mm^2^. Remaining mice were euthanized and spleens were harvested when more than half of the mice in a group reached tumor endpoint. *Randomization:* Prior to therapy, mice were randomized based on tumor size. *Blinding:* Tumors were measured using L × W measurements via calipers by personnel blinded to treatment group.

### Statistical Analysis

Tumor area results were transformed using the natural logarithm for data analysis. Mixed effects linear regression models with a random component to account for the correlation of the repeated measure within a mouse were used to estimate tumor area over time. In circumstances where linearity assumptions were not met, polynomial regression models were used (*27*). Linear combinations of the resulting model coefficients were used to construct estimates for the slope differences with 95% confidence intervals where applicable. For polynomial models, estimates were constructed for the differences in area between groups on the last day where at least one mouse was alive in all groups. Experiments with multiple groups were analyzed using one-way analysis of variance (ANOVA) with post comparison of all pair wise groups using Tukey’s range test. Experiments comparing two groups were analyzed using a Student’s t test. The center values are the mean and error bars are calculated as the SEM. TCRβ sequencing analysis was based on the log-linear model and the ‘relative risks’ calculated with a 95% confidence interval.

### Subset Isolation

De-identified, normal human donor peripheral blood cells were purchased as a buffy coat (Plasma Consultants) or leukapheresis (Research Blood Components). PBL were enriched using Lymphocyte Separation Media (Mediatech). CD4^+^ T cells were negatively isolated using magnetic bead separation (Dynabeads, Invitrogen) and plated in culture medium with a low concentration of rhIL-2 (20 IU/ml; NIH repository) overnight. For *in vivo* studies, CD8^+^ T cells were positively isolated prior to the enrichment of CD4^+^ T cells. The following morning CD4^+^ T cells were stained using PE-CD26 (C5A5b), AlexaFluor647-CXCR3 (G025H7), PECy7-CCR6 (G034E3, Biolegend), FITC-CCR4 (205410, R&D Systems) and APCCy7-CD4 (OKT4, BD Pharmingen). Cells were sorted based on the following gating strategies: bulk CD4: CD4^+^; Th1: CD4^+^CCR6^−^CCR4^−^CXCR3^+^; Th2: CD4^+^CCR6^−^CCR4^+^CXCR3^−^; Th17: CD4^+^CCR6^+^CCR4^+^; CD26: CD4^+^CD26^high^. Cells were sorted on a BD FACSAria IIu Cell Sorter or on a Beckman MoFlo Astrios High Speed Cell Sorter.

### T cell culture

T cell subsets were expanded in RPMI 1640 culture medium supplemented with non-essential amino acids, L-glutamine, sodium pyruvate, HEPES, Pen/Strep, β-mercaptoethanol and FBS. Cells were cultured at either a 1:1 or 1:10 bead to T cell ratio. Magnetic beads (Dynabeads, Life Technologies) coated with antibodies to CD3 (OKT3) and/or ICOS (ISA-3, eBioscience) were produced in the lab according to manufacturers’ protocols. One hundred IU/ml rhIL-2 (NIH repository) was added on day 2 and media was replaced as needed.

### T cell transduction

To generate mesothelin-specific T cells, αCD3/ICOS-activated, sorted CD4^+^ and bulk CD8^+^ T cells were transduced with a chimeric anti-mesothelin single-chain variable fragment (scFv) fusion protein containing the T cell receptor ζ (TCRζ) signaling domain (1^st^-gen-Meso-CAR) or a truncated CD3ζ non-signaling domain (Δζ) that was generated as described previously (*9*). CAR expression was determined using a flow cytometry antibody specific for the murine F(ab’)_2_ fragment (Jackson ImmunoResearch, 115-606-006).

### Flow cytometry

For intracellular staining data, cells were stimulated with PMA/Ionomycin. After one hour, Monensin (Biolegend) was added and incubated for another 3 hours. Following surface staining, intracellular staining with antibodies was performed according to the manufacturer’s protocol using Fix and Perm buffers (Biolegend). Data were acquired on a BD FACSVerse or LSRII X-20 (BD Biosciences) and analyzed using FlowJo software (Tree Star, Ashland, OR).

### MicroArray

RNA was isolated from sorted CD4^+^ T cells using the Qiagen RNeasy Mini kit and frozen. RNA was submitted to Phalanx Biotech Group for processing on their OneArray platform (San Diego, CA). *Heatmap and PCA clustering:* Graphing was performed in R (version 3.1.2) using gplots (version 2.16.0). Log_2_ values for CD4^+^ cells were averaged and used as baseline for the genes of interest. For each individual sample the fold change relative to baseline was calculated and the median value for the triplicates was calculated and used for generating figures.

### ATAC sequencing

Sorted CD4^+^ T cells were cryopreserved in CryoStor and sent for analysis. Naïve cells were sorted based on expression of CCR7 and CD45RA. ATAC-seq was performed by Epinomics according to the protocol described by Buenrostro et al. (*28*). Fifty thousand sorted T cells were frozen using Cryostor CS10 freeze media (BioLife Solutions) and shipped on dry ice for processing and analysis to Epinomics (Menlo Park, CA).

### T Cell Receptor β sequencing

Sorted T cells were centrifuged and washed in PBS, and genomic DNA was extracted using Wizard Genomic DNA purification kit (Promega). The quantity and purity of genomic DNA was assessed through spectrophotometric analysis using NanoDrop (ThermoScientific). Amplification of TCR genes was done within the lab using the ImmunoSEQ hsTCRβ kit (Adaptive Biotechnologies Corp., Seattle, WA) according to the manual. Survey sequencing of TCRβ was performed by the Hollings Cancer Center Genomics Core using the Illumina MiSeq platform.

### Single cell RNA sequencing

Sorted Th17, CD26^high^ T cells or bulk CD4^+^ T cells were cryopreserved and sent for analysis to David H. Murdock Medical Research Institutes (DHMRI) Genomics core. Genomic DNA was analyzed using the Chromium Controller instrument (10X Genomics, Pleasanton, CA) which utilizes molecular barcoding to generate single cell transcriptome data (*29*). Sequencing of the prepared samples was performed with a HiSeq2500 platform (Illumina). Data were analyzed with Long Ranger and visualized with Loupe (10X Genomics).

### *In vitro* cytotoxicity assay

Sorted CD4^+^ T cell subsets were activated with αCD3/ICOS beads and engineered to be mesothelin-specific using a lentiviral CAR. Following a 10-day expansion, equal numbers of transduced T cells were co-cultured overnight with target cells. For the CAR, meso-expressing K562 cells (pre-stained with Cell Trace Violet, Molecular Probes) serially diluted in the presence of CD107A (Pharmingen). K562 lysis was determined by 7-AAD (Pharmingen) uptake. K562-meso cells were tested for mycoplasma (MycoAlert, Lonza) and mesothelin (R&D Systems, FAB32652) expression during expansion.

### Mice and tumor line

NOD SCID gamma chain knockout mice (NSG, The Jackson Laboratory) were bred at the University of Pennsylvania or at the Medical University of South Carolina. NSG mice were given ad libitum access to autoclaved food and acidified water. M108 xenograft tumors (gift C.H. June), described previously (*9*), were tested for mycoplasma during expansion (Lonza).

#### Ethics Approval

Human peripheral blood was not collected specifically for the purposes of this research and all samples were distributed to the lab in a deidentified manner. Therefore, this portion of our research was not subject to IRB oversight. All animal studies were approved by the Institutional Animal Care and Use Committee (IACUC) at the Medical University of South Carolina.

## General

The authors thank Leah Stefanik, Kristina Schwartz and Marshall Diven of the Department of Microbiology & Immunology for their assistance. We thank Adam Soloff and Zachary McPherson at the MUSC Flow Cytometry and Cell Sorting Core and Chris Fuchs at Regenerative Medicine Department Flow Core for sorting cells. We also thank Dimitrios Arhontoulis, Ephraim Ansa-Addo, Krishnamurthy Thyagarajan, and Juan Varela for critical reading and feedback. We want to acknowledge Elizabeth Garrett-Mayer for TCRβ data analysis, Arman Aksoy and Jeff Martello for assistance with the single cell analysis, and Epinomics for collaboration and ATAC-seq analysis. Finally, we want to thank Carl June (University of Pennsylvania) and Nicholas Restifo (Surgery Branch, NCI) for reagents and support.

## Author contributions

M.H.N. designed and executed experiments, analyzed the data, created the figures, and wrote and edited the manuscript; H.M.K performed experiments, analyzed data, created figures, wrote and edited the manuscript; S.R.B, J.S.B., L.W.H., K.M., and M.M.W performed experiments; K.E.A., P.G., and M.J.Z analyzed the data, S.M., M.P.R., M.I.N., H.E.B designed experiments and edited the manuscript, and C.M.P. directed the project, designed experiments, and edited the manuscript. All authors critically read and approved the manuscript.

## Data and materials availability

For original data, please contact. Microarray data can be found at GEO accession number GSE106726.

## Supplementary Figures

**Figure S1.**
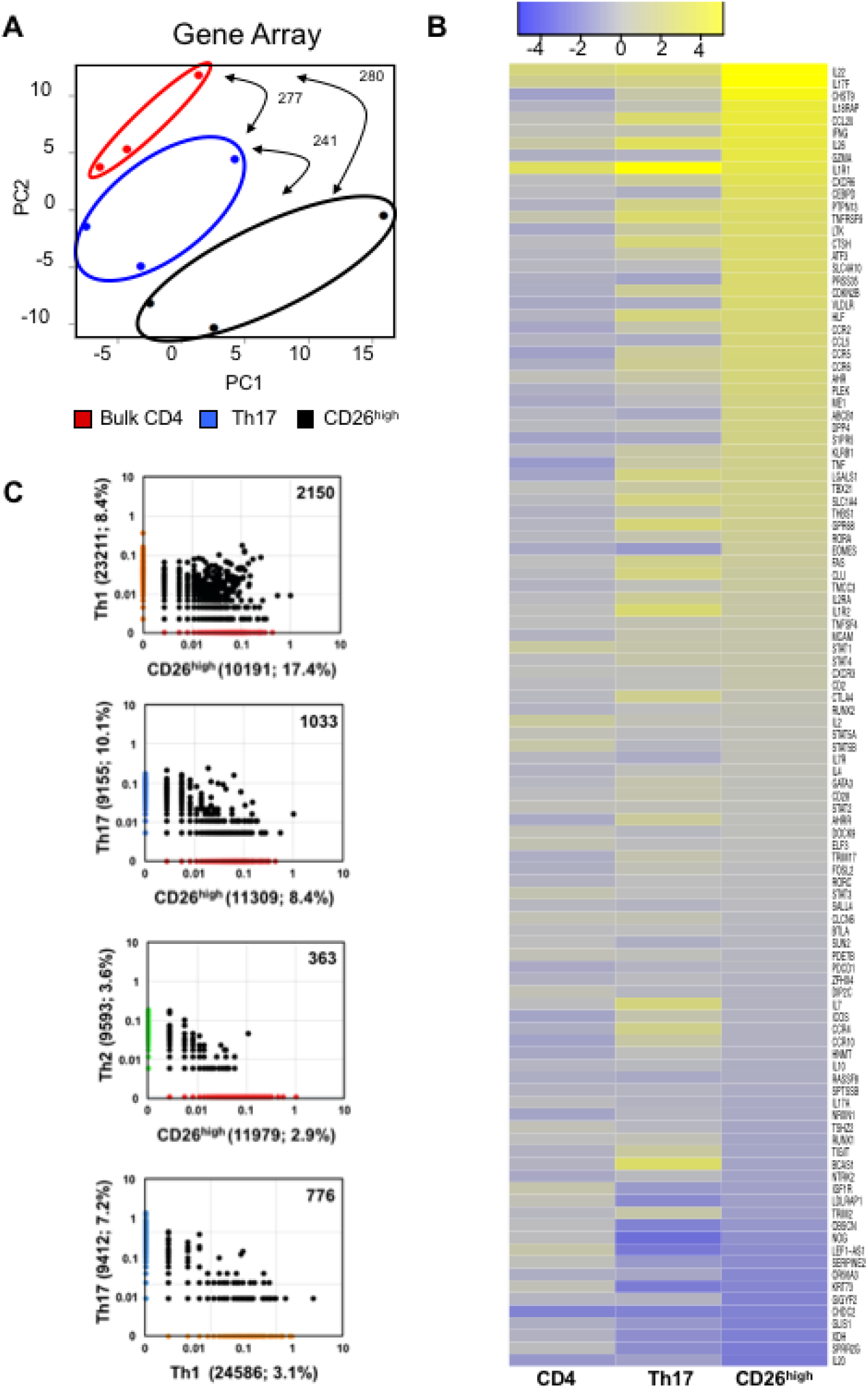
CD26^high^ T cells have a unique molecular phenotype and TCRβ repertoire from classic helper cells. **A-B)** RNA was isolated from 3 normal donors’ sorted T cell subsets and gene expression levels were determined by OneArray on day 0. **A)** Principal component analysis and **B)** Heat map of log2-fold change in expression of genes with the highest or lowest expression in CD26^high^ T cells. **C)** T cell subsets were sorted from peripheral blood of normal human donors based on surface chemokine receptor expression (Th1 (CXCR3^+^CCR6^−^), Th2 (CCR4+CXCR3-) Th17 (CCR4^+^CCR6^+^), CD26^high^ (top 5%)). DNA was isolated, TCRβ sequences were expanded using an immunoSEQ kit and subsequently sequenced. Data shown is the graphical representation of TCR overlap between indicated T helper subsets. Representative from 4 donors.

**Figure S2.**
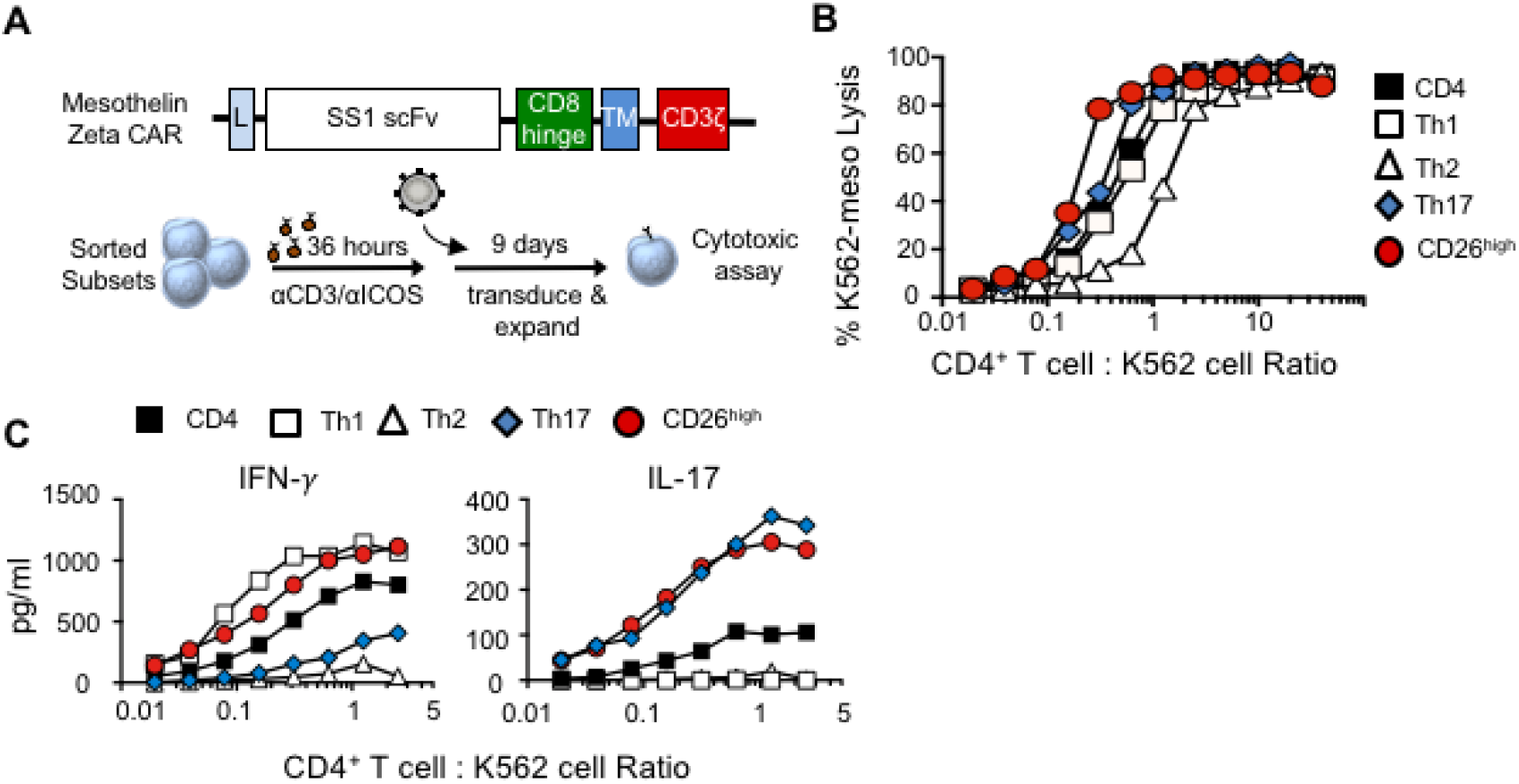
CD26^high^ cells are cytotoxic and polyfunctional *in vitro* when engineered with a chimeric antigen receptor. **A**) Transduction method. αCD3/ICOS-stimulated CD4^+^ T cell subsets were genetically engineered with a 1^st^ generation mesothelin-specific CAR. Cells were expanded for 6 days and analyzed by flow cytometry for CAR expression prior to use. **B**) Percentage of K562-meso cells that were lysed by effector CD4^+^ T cell subsets. **C**) Cytokine secretion determined by ELISA. Representative of 3 experiments.

**Figure S3:**
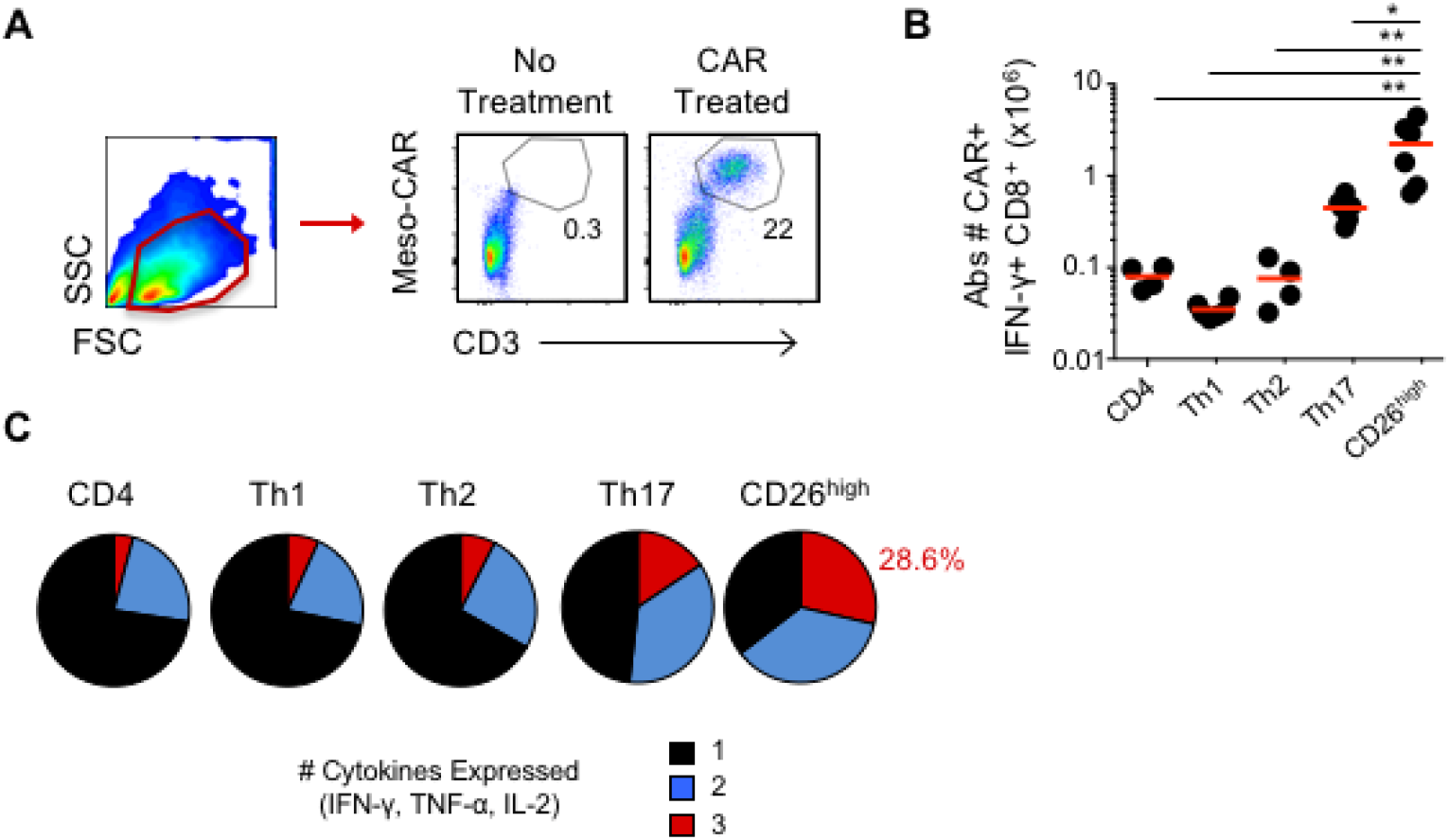
Human CD26^high^ T cells improve function of co-transferred CD8^+^ meso-CAR T cells. **A-C)** Sorted Th1, Th2 Th17, CD26^high^ or CD4^+^ cells were transferred into mesothelioma-bearing NSG mice as described in Figure 3D. **A)** Representative flow cytometry gating for CAR T *in vivo*. **B)** Total number of splenic IFN-γ-producing CD8^+^meso-CAR^+^ cells. n=4-6 mice/group. Compared to CD26^high^ *, *P* < 0.05; **, *P* < 0.01; ANOVA, Tukey post-hoc comparisons. **C)** Simultaneous intracellular cytokine production in spleen CAR^+^CD8^+^ cells. Average of 4-6 mice/group.

**Figure S4.**
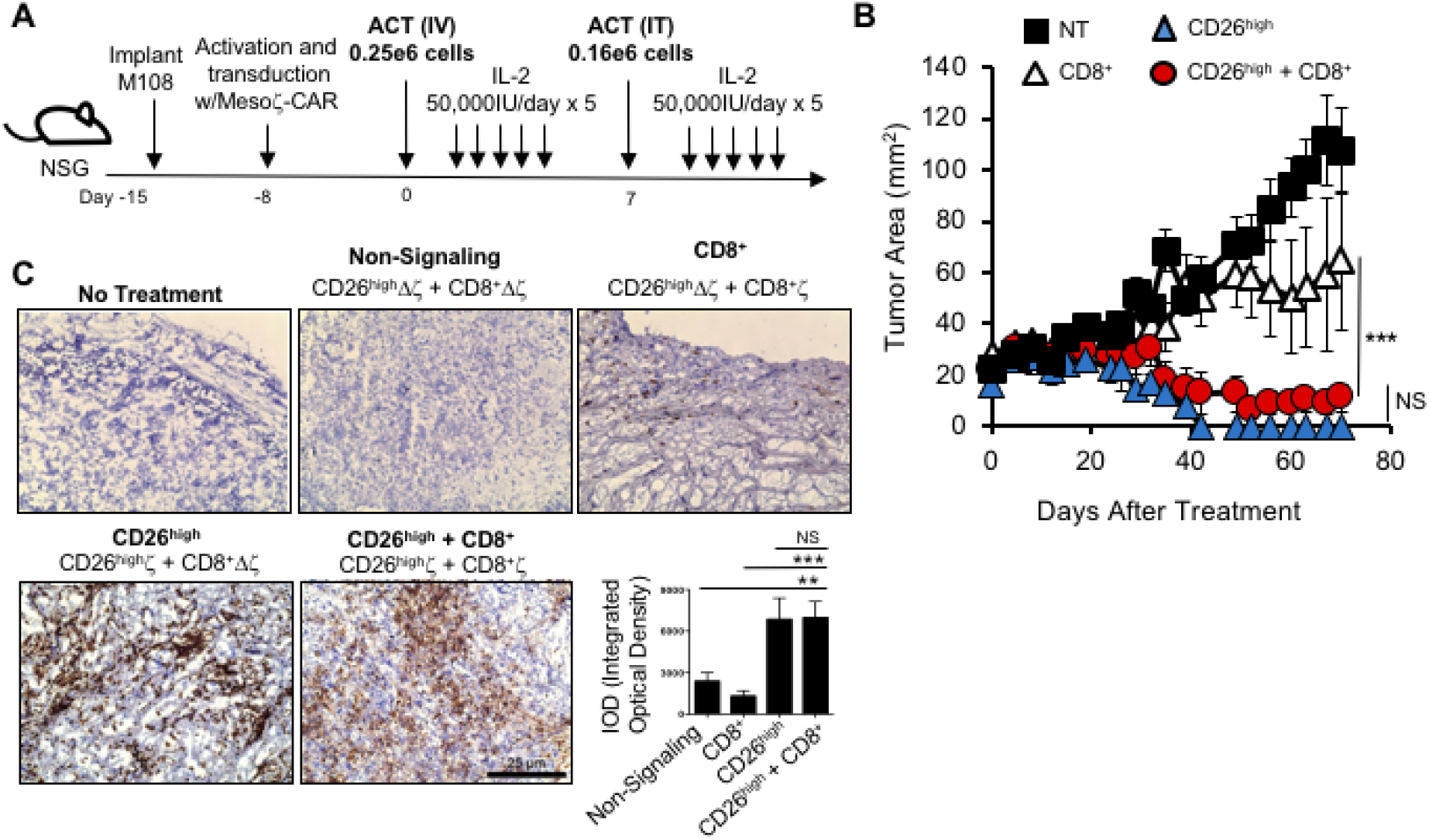
Human CD26^high^ meso-CAR T cells do not require CD8^+^ CAR T cells for antitumor responses. **A-B)** Mesothelioma-bearing NSG mice were treated with CD26^high^ T cells co-infused with or without CD8^+^ T cells all engineered with 1^st^-gen-meso-CAR. Two infusions of cells were given one week apart (250,000 cells i.v.; 160,000 cells i.t.). 5-6 mice/group. All groups were significantly different, *P* < 0.001, except CD8^+^ + CD26^high^ vs. CD26^high^, *P* < 0.43. **C)** Immunohistochemistry staining of M108 from NSG mice treated with 1.45×10^6^ CD26^high^ and CD8^+^ T cells transduced with a 1^st^-gen-Meso-CAR having either full-length CD3ζ signaling or a non-signaling truncated version (∆ζ). Staining of human CD45 and hematoxylin on day 84 post-transfer (x10; 3 or 4 mice/group; average IOD from 10 images). Compared to CD26^high^ **, *P* < 0.01; ***, *P* < 0.001; ANOVA.

